# Grid Codes versus Multi-Scale, Multi-Field Place Codes for Space

**DOI:** 10.1101/2023.06.18.545252

**Authors:** Robin Dietrich, Nicolai Waniek, Martin Stemmler, Alois Knoll

**Author notes:** Correspondence: Robin Dietrich.

## Abstract

Recent work on bats flying over long distances has revealed that single hippocampal cells represent space on many scales, suggesting that these cells simultaneously participate in multiple neuronal networks to yield a multi-scale, multi-field place cell code. While the first theoretical analyses revealed that this code outperforms classical single-scale, single-field place codes, it remains an open question what the performance boundaries of this code are, what functional properties the network responsible for this code has, and how it compares to a highly regular grid code, in which cells form distinct modules, each with its own attractor dynamics on the network.

In this paper we address these questions with rigorous analyses of comprehensive simulations. Specifically, we perform an evolutionary optimization of several multi-scale, multi-field place cell networks and compare the results against a single-scale, single-field as well as against a simple grid code. We focus on two main characteristics: the general performance of the code itself and the dynamics of the network generating it. Our simulation experiments show that, under normal conditions, the grid code easily outperforms any multi-scale, multi-field place code with respect to decoding accuracy. However, the latter is more robust to noise and lesions, such as drop-out. The robustness comes at a cost, as the grid code requires a significantly smaller number of neurons and fields per neuron. Further analyses of the network dynamics also revealed that the proposed topology of multi-scale, multi-field place cells does not, in fact, result in a continuous attractor network. More precisely, the simulated networks do not maintain activity bumps without position specific input. The multi-scale, multi-field code, therefore, seems to be a compromise between a place code and a grid code that invokes a trade-off between accurate positional encoding and robustness.

## 1 INTRODUCTION

Navigating large and complex environments is a non-trivial task. It requires perception of the environment, a subsequent map formed by these perceptions, a localization mechanism within it as well as a method for navigating between two points in the map Thrun et al. (2005). Humans, as well as mammals, in general are able to perform this task seamlessly, whether in a small room or a large environment, such as a city. The neural formations responsible for the respective tasks have been investigated for decades. Yet, the exact representation a mammal keeps of an environment remains covert.

The hippocampal formation has been identified as a primary unit for the computation and storage of a neuronal spatial map akin the cognitive map theory by Tolman (1948) ever since the discovery of place cells (PCs) by O’Keefe and Dostrovsky (1971). PCs were found in the CA1 and CA3 sub-regions of the Hippocampus and commonly show singular or only few prominent areas of maximal firing activity relative to the environment in which an animal is located, the cells’ so-called place fields. This led to the – nowadays widely accepted – hypothesis that these neurons discretize a continuous environment into a finite number of place fields. In turn, this motivated a plethora of biological experiments as well as modelling approaches, covering a wide range of aspects, including the influence on the firing field size/shape caused by different factors, such as the environment O’Keefe and Burgess (1996), the animal speed Ahmed and Mehta (2012) or the recording location within the hippocampus O’Keefe and Burgess (1996). These studies revealed that place cells can express multiple place fields under certain circumstances Kjelstrup et al. (2008); Park et al. (2011); Davidson et al. (2009); Rich et al. (2014) and that the size of these fields can vary O’Keefe and Burgess (1996); Fenton et al. (2008). The majority of these experiments were, however, conducted in small, confined spaces, since the technology and hardware that is required for neural recordings did not support large and unconfined environments as of the time of the studies.

The advancement of hippocampal recording technology towards wireless communication recently allowed to conduct experiments in large scale environments and to study different firing properties of place cells (PCs) in dorsal CA1 of the hippocampus in such surroundings Eliav et al. (2021a); Harland et al. (2021). Both studies reported place cells with multiple, differently sized place fields - a *multi-scale multi-field (MSMF) place code*. This code is similar to the *grid code* produced by grid cells found in the medial entorhinal cortex (MEC) Hafting et al. (2005). While each grid cell also maintains multiple fields, the size of these fields is constant per neuron and only changes across so called modules of neurons with the same scale. The fields are distributed regularly in a hexagonal pattern forming an optimal code for arbitrary spaces Mathis et al. (2015). In contrast to that, the experiments performed by Eliav et al. (2021a) revealed the MSMF code for neurons in the hippocampus of bats flying through a 1-dimensional, 200*m* long tunnel. Harland et al. (2021) identified this property of PCs in rats foraging within a 2-dimensional, 18, 6*m*^2^ open arena. Eliav et al. (2021a) also performed a theoretical analysis of the concept to demonstrate the effectiveness of this multi-scale code compared to single-scale. The authors show, that in order to achieve a localization error of *<* 2*m*, a single-field model requires more than 20 times as many neurons than an MSMF model. This analysis further shows, that using a fixed number of 50 neurons, the decoding error is 100 times better with the MSMF model than with the single-field model.

Beyond this theoretical analysis, Eliav et al. (2021a) also introduce two neuronal models in a computational analysis, which could explain this MSMF code - a continuous attractor network (CAN) and a feedforward model receiving input from either CA3 place cells or MEC grid cells. The 1D CAN consists of multiple, distinct, differently sized, overlapping attractor networks, each of which containing the same amount of neurons, as shown in Fig. 1. The authors perform experiments in a 200*m* long environment using 4000 neurons (1200 neurons per attractor) in order to demonstrate that this network is capable of generating a MSMF code. The analysis of this model beyond that is quite limited. The field sizes were analyzed, as shown in Fig. 1, but there were no experiments conducted for evaluating the decoding accuracy of said network.

**Figure 1.**
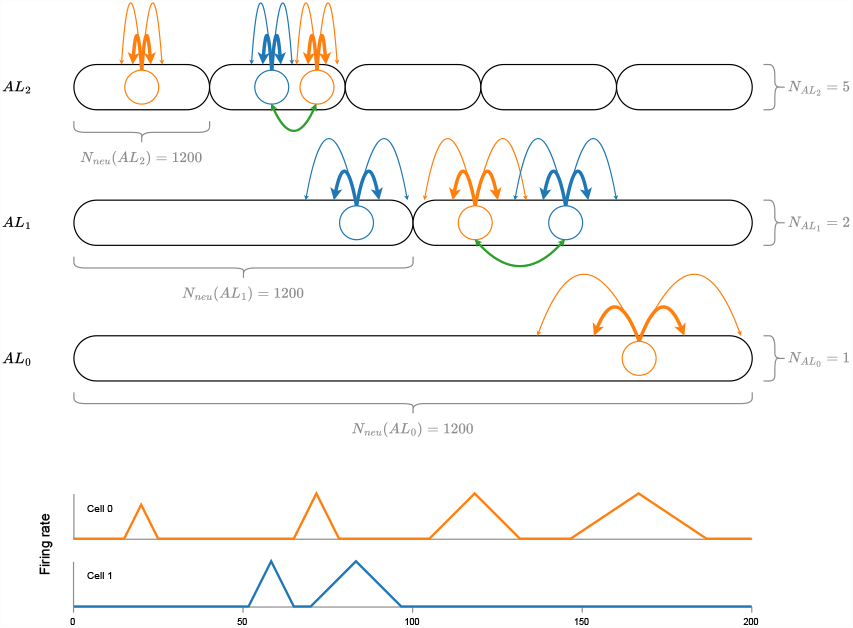
A visualization of the CAN model introduced by Eliav et al. (2021a) with a total of 8 attractor networks, coupled together by neurons in the same attractor (green lines). Each attractor network consists of the same amount of neurons (*N*_*neu*_ = 1200), drawn randomly from a total number of 4000 neurons. At the bottom of the figure, an idealized firing rate for the two neurons (blue, orange) is shown. Note, that although the size of a firing field is generally pre-determined by the respective attractor network, it can vary depending on the overall connectivity of the neuron. See the first two fields of cell 0 for an example.

The results from the theoretical and computational analysis of the MSMF code performed by Eliav et al. (2021a) suggest the discovery of a superior coding scheme for the position of an animal. Yet, these results raise several important questions, both, from a neuroscientific and a computational point of view. First, it has been shown previously, that the ”traditional” single-scale, single-field place code is outperformed by the grid code Mathis et al. (2012) and that the grid code also maintains an optimal distribution of fields per neuron for arbitrary spaces Mathis et al. (2015). This raises the question, whether this MSMF code has any advantages over the grid code with respect to the decoding accuracy, energy consumption or robustness. Second, the discrepancy between the number of neurons used for the theoretical (50) as well as the computational analysis (4000) by Eliav et al. (2021a) is non-negligible and opens up the question, whether a network with a computationally realistic neuron model as well as interconnections would be able to achieve such a performance. Can an optimization algorithm find a network with an accuracy close to the one from the theoretical experiments? How would the neurons have to be connected? What would an optimal distribution of the fields look like? Finally, when inspecting the general structure of the original MSMF network in combination with the distribution of the fields in the experiments, one naturally wonders about the dynamics of a network for such a code. How do the coupled attractors in the MSMF network interact and interfer with each other? Would this still be a continuous or rather a discrete attractor network?

We will answer these questions in the remainder of this paper. Our approach is to optimize several candidate networks using evolutionary optimization techniques and compare their performances in different scenarios. The main contributions of our work are as follows:

- we perform an in depth analysis of the MSMF CAN model proposed by Eliav et al. (2021a). While we can confirm many of their findings, we find further insights and significant properties,
- we optimize the parameters for both, the attractor network introduced by Eliav et al. (2021a) as well as a model based on their theoretical evaluation, to determine the optimal set of parameters for each,
- we demonstrate that while these models do work with mixed field sizes, they achieve a much better decoding accuracy when constructed of many small fields instead of a variety of field sizes. This is in strong contrast to the theoretical analysis reported in Eliav et al. (2021a),
- we show that MSMF models are easily outperformed by a simple grid code. This raises concerns about the likelihood of such codes being used in the mammalian hippocampus for exact position encoding of an animal,
- we demonstrate that MSMF models are significantly more robust against noise compared to grid as well as single field models,
- we show that lateral connections in both MSMF models do not form the basis of an actual CAN but they do improve the decoding accuracy under specific circumstances,
- finally, we provide an openly accessible framework for easily optimizing and evaluating the different networks.

## 2 METHODS

### 2.1 Network Models

In the following we will introduce the different MSMF models evaluated throughout the paper, together with the grid and single field model used as a baseline. An overview of each network’s parameters is given in Table S9. The dynamics and neuron models are identical for all networks and will be described in section 2.2.

#### 2.1.1 Single-Scale Single-Field Model

As a baseline, we implemented a simple single-scale single-field (SSSF) model, based on the F-MSMF model but with only one attractor. The neurons within this attractor are distributed uniformly over the entire environment.

#### 2.1.2 Grid Cell Model

In order to compare the MSMF place cell models to an optimal encoding of space, we also implemented a one-dimensional version of a grid model without lateral connections. Analogous to 2D grid models, this model consists of multiple modules *N*_*mod*_, each containing a fixed number of neurons 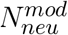. Each module further has a certain scale, starting with the minimum defined scale 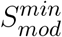 and increasing per module by the module scale factor *S*_*mod*_. The neurons within each model then maintain regularly recurring firing fields based on this scale and a certain, increasing offset for each neuron within a module, generating a 1D grid code.

#### 2.1.3 Fixed Multi-Scale Multi-Field Model

The first MSMF model we consider is adapted from Eliav et al. (2021a). The authors introduce a network for 1D environments, where the neurons are organized not just in a single line attractor, but in multiple, differently sized line attractors interacting with each other. We call this a *fixed* MSMF network (F-MSMF), due to the fixed, predetermined number of line attractors. A schematic of this architecture is visualized in Fig. 1. The network consists of multiple, distinct attractor networks, distributed over three different stages, each having a different interaction length *L*_*int*_, resulting in individual field sizes per attractor.

This way of organizing and scaling the attractors leads to different field sizes on the different levels, as the overall attractor size changes while the number of neurons per attractor is constant. In the model by Eliav et al. (2021a), a pool of neurons *N*_*neu*_ = 4000 is created at the beginning and each of these neurons can participate in each of the attractors. The probability of participation of one neuron in any one attractor is set by Eliav et al. to *P*_*att*_ = 0.3 leading to a total number of 1200 neurons/fields per attractor. Since the attractors span over different lengths of the environment while maintaining the same number of neurons, this leads to the different field sizes per attractor. The result is a multi-scale, multi-field code, as each neuron can be part of not one but multiple attractors (multi-field) with different field sizes per attractor (multi-scale). While Eliav et al. (2021a) do perform some general analysis of this model (field sizes, distribution) they do not investigate the performance (positional decoding accuracy) or efficiency (potential energy consumption, number of neurons) of the network as they did in the theoretical analysis described before.

The default parameters used in our experiments for the field and attractor generation of this model are listed in Table S9. Most of these parameters are identical with the ones from the evaluations by Eliav et al. (2021a). It is unclear if the parameters reported by Eliav et al. (2021a) were selected to stabilize the network, or if they were extracted from real world recordings. For further details regarding this model, we refer the reader to Eliav et al. (2021a,b).

#### 2.1.4 Dynamic Multi-Scale Multi-Field Model

Based on the insights from Eliav et al. (2021a), we have developed a new *dynamic* MSMF model (D-MSMF) composed of a dynamic number of attractor networks. The model has the general architecture of a CAN but dos not fully comply with all properties of either a continuous or a discrete attractor network, settling it somewhere in between. The core idea behind this approach is, that only connections between two neurons with similar field sizes should be modeled and incorporated. This generalizes the idea of having multiple, interacting attractors proposed by Eliav et al. (2021a). The authors created only three different categories of sizes of fields, or attractors. These fields further spanned the whole environment uniformly.

A visualization of a few neurons, together with their fields and respective connections, taken from a D-MSMF network, are shown in Fig. 2a. In order to generate such a network, we first create a population of *N*_*neu*_ neurons and then sample fields for each of the neurons, using the same gamma distribution as Eliav et al. (2021a) for their theoretical analysis. We base the field distribution on these results, as they are in turn based on the measured and calculated values from the biological experiments. New fields for a neuron are then generated until the overall size Σ_*fs*_ of all fields of a neuron *n* reaches a certain threshold 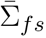. This threshold is defined using values retrieved from the biological experiments as well Eliav et al. (2021a).

**Figure 2.**
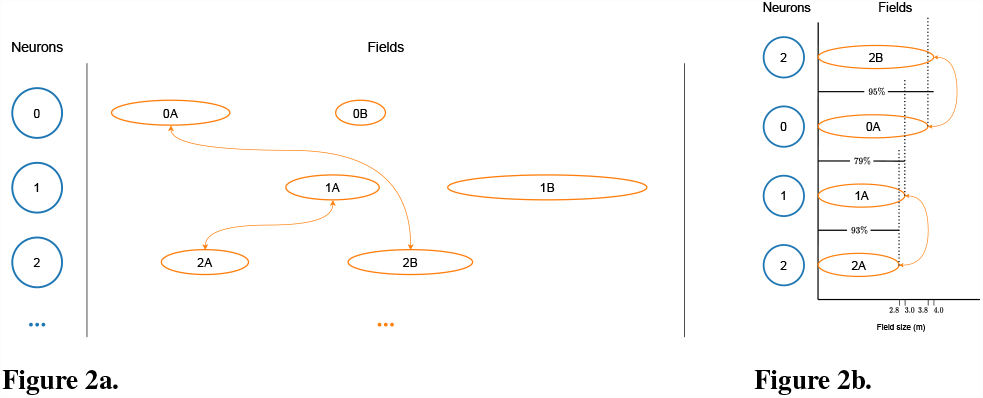
Visualizations of the MSMF model developed by us based on the theoretical model from Eliav et al. (2021a). **(a)** The differently sized firing fields of three neurons. Only connections between neurons with fields of similar size (0A↔2B, 1A↔2A) are modeled. **(b)** The size difference between the firing fields, shown in detail. In this example a threshold *TH*_*fs*_ = 0.9 = 90% was selected.

Subsequently, the connection weights between all neurons are calculated. For this purpose, we define a threshold *TH*_*fsr*_ for the ratio between the size of two fields. We then compare the sizes of all fields of two neurons (*n*_0_, *n*_1_). The overall connection strength between these two neurons is generally defined by the distance between all fields of these neurons. In order to achieve a similar architecture as Eliav et al. (2021a) with their CAN model, we do, however, only take those fields into account, whose ratio is above the threshold *TH*_*fsr*_, i.e.

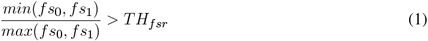

for fields with sizes *fs*_0_ ∈ *n*_0_ and *fs*_1_ ∈ *n*_1_. A simplified diagram of this mechanism for connection weight calculation in an MSMF network is visualized in Fig. 2b. Here a threshold of *TH*_*fsr*_ = 0.8, was chosen, hence only two connections between the three depicted neurons will be created. The first synapse connects neurons *n*_0_ and *n*_2_ with a weight based on fields *f*_0*A*_ and *f*_2*B*_. The second synapse connects neurons *n*_1_ and *n*_2_ with a weight based on fields *f*_1*A*_ and *f*_2*A*_. This connection scheme is a generalization of the F-MSMF model, since that model also creates multiple connections between two neurons based on the interaction of the respective neurons in the respective attractors. If two neurons have a connection in the F-MSMF model, then they also have two fields of similar size, since they participate in the same attractor. We use the D-MSMF model now in order to further investigate the influence of the field size on the connection probability between two neurons. This is possible, since the connection probability can be easily adjusted with the *TH*_*fsr*_ parameter. The major difference between the two models is the distribution of the field sizes and the fact, that in the F-MSMF model all attractors span uniformly over (a part of) the environment. In the D-MSMF model, this is not necessarily the case. Due to the dynamic creation of fields and attractors, the position of a field within an attractor is not predetermined.

The parameters used for generating the fields are listed in Table S9. The dynamics of the network are the same as for the F-MSMF network. They are described in section 2.2.

### 2.2 Neuron Model

The dynamics of all networks introduced in Section 2.1 are based on the ones defined by Eliav et al. in Eliav et al. (2021b). Here, we therefore only summarize the essentials to understand the remainder of this work while complementing it with our own additions.

According to Eliav et al. (2021b), the dynamics for a single neuron *i* are defined by

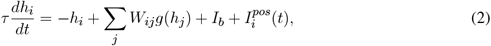

where *W*_*ij*_ defines the overall weight between neuron *i* and neuron *j* according to the chosen model and *I*_*b*_*ck* defines a uniform background input (noise). The neuronal gain function (*g*(*h*)) has a threshold-linear form as follows

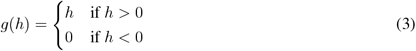

The positional input 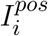 on the other hand defines the individual input each neuron receives based on the position of its fields and the respective distance of those to the current position of the agent

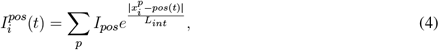

where *pos*(*t*) defines the position of the agent at time *t* within the 1D environment, assuming a constant speed of 10*m/s*.

Beyond these general dynamics of the networks, we also introduced a variable, noisy background input, replacing *I*_*bck*_ in some experiments. The noisy background input is defined by a mean 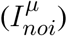 as well as a standard deviation 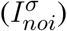 of the normal distribution generating the noisy input values.

### 2.3 Optimization

As one of the most prevalent biologically inspired optimization methods, *evolutionary optimization* is a prime candidate for finding the most suitable parameter configurations for the models defined in this section. Specifically, because it allows the optimization without prior knowledge and hence limiting the number of assumptions that need to be made. Within this section we will briefly discuss how we used evolutionary optimization for finding new parameter configurations, leading to an improved accuracy or energy efficiency consumption of the models.

The process of our algorithm is depicted in Figure 3 and based on Simon (2013). We first *generate* a set of entities (commonly *N*_*pop*_ = 20 with some selected network parameters being randomly initialized. For this random initialization, we defined a lower as well as an upper bound of the values for each parameter individually. Additionally, we only allowed values based on a certain step size, which was either an integer or a floating point number, based on the type of the parameter. This reduces the search space from the beginning and allows to run experiments faster. As a second step, the networks of all entities are *evaluated*, this commonly encompasses 20 runs of the same network. This ensures, that the calculated metrics are representative, as the decoded positional accuracy can highly diverge for the same network parameters but different field locations for the neurons. We elaborate more on this within our results and discussion. The fitness function we use is based on the mean or median error of the network and is defined as follows

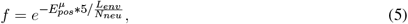

where 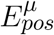 defines the mean of multiple mean positional decoding errors, calculated from several runs with the same network parameters, *L*_*env*_ is the total length of the environment in meters and *N*_*neu*_ is the total number of neurons. The constant 5 was simply introduced to scale the fitness function a bit up. Subsequently, a number of entities to keep for the next generation is *selected* from the entire population. This is done using fitness-weighting, i.e. the entities are ordered by their fitness first and then a subset of them is selected based on the defined selection rate *R*_*sel*_ (commonly *R*_*sel*_ = 0.2). From this new set of entities, parents are chosen for *mating*, with a probability proportional to their fitness. Based on two chosen parents, a child entity is generated with parameters inherited from both parents. This inheritance is performed randomly. An integer is randomly generated, dividing the number of optimization parameters in two halves, one from each parent. The optimization parameters of the children created in this step are then randomly mutated with a probability *P*_*mut*_ (commonly *P*_*mut*_ = 0.2). As a final step a new population is created from the children. In all of our experiments, we additionally kept the entity with the best fitness from the selected entities constant without mating or mutating its parameters.

**Figure 3.**
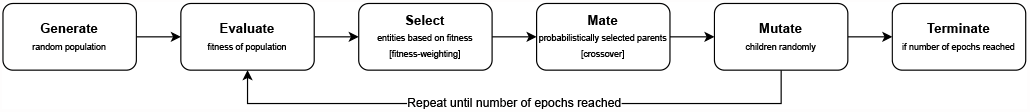
A visualization of the evolutionary optimization process.

This process is continued until the defined number of epochs is reached (commonly *EP* = 3000).

## 3 EXPERIMENTAL EVALUATION

The models introduced in the previous section form the basis of our simulated experiments presented within this section. We first describe the general setup of the experiments. Then we introduce the results of the baseline models by Eliav et al. (2021a), as well as their optimization with and without lateral connections. With these evaluations we demonstrate the usefulness of the MSMF code itself as well as possible network structures for generating them.

### 3.1 Experimental Setup and Metrics

In order to rule out outliers, each experiment presented in this section with a single set of parameters was evaluated by performing 20 simulations of the same network and calculating the statistics (mean, median, standard deviation) of the positional error, the number of fields and other metrics. We commonly make use of the median, since the distribution of most metrics over the 20 runs are not Gaussian.

For some of the evaluations we also use an efficiency measurement as a comparison metric. We therefore define the median expected energy consumption for multiple runs of the same network as

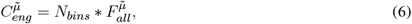

where *N*_*bins*_ is the total number of bins of the environment (for most experiments 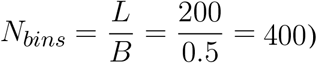 and 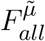 is the mean in-field activity (firing-rate) of all fields (active as well as inactive).

The original models as well as the optimized ones will be abbreviated by *F/D/G-Org* and *F/D/G-Opt*, respectively. The first letter indicates the type of model, i.e. F-MSMF (*F*), D-MSMF (*D*) or grid (*G*). We indicate that a model contains lateral connections (*D-Org-1*^+^) or not (*D-Org-1*^*−*^), and also whether the connections in this model were optimized (*D-Org-1*^+*o*^). In case the model receives a uniform background input (*I*_*bck*_), this is indicated by a subscript “*β*” 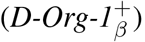.

### 3.2 Original Models

The first part of our evaluations consists of experiments performed with the original models and simulations introduced by Eliav et al. (2021a). We evaluated both, the F-MSMF and D-MSMF model, in order to analyse their positional encoding performance and answer the question, whether these models are generally capable of reproducing the results of the theoretical analysis by Eliav et al. (2021a) and identifying potential improvement possibilities.

In our first experiment, we evaluated the F-MSMF model with identical parameters as proposed by Eliav et al. (2021a), i.e. we simulated the network with a total number of *N*_*neu*_ = 4000 neurons. We then modified the parameters of the lateral connections in the network (*W*_*exc*_, *W*_*inh*_) as well as the noise or background input (*I*_*bck*_) in order to evaluate their impact on the encoding performance of the network. The statistics of the mean positional error for four models with different parameter combinations are visualized in Fig. 4a. This simulation shows, that all three parameters have a significant influence on the accuracy of the network. Setting the background input as well as all lateral connections to zero results in a decrease of the median of the average positional error 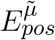 by 1.128*m* (1.226*m →−* 0.098*m*). Especially the background input has a significantly negative effect on the median performance (see models 3, 4). The lateral connections, on the other hand, seem to have a strong influence on the standard deviation, leading to a broader overall distribution including both, networks with better as well as worse performances than without lateral connections. These results are further backed by the same experiment performed with only *N*_*neu*_ = 50 neurons, shown in Fig. 4b. The number of neurons was set to 50 here because this is the same number of neurons that was used by Eliav et al. (2021a) in their theoretical evaluations.

**Figure 4.**
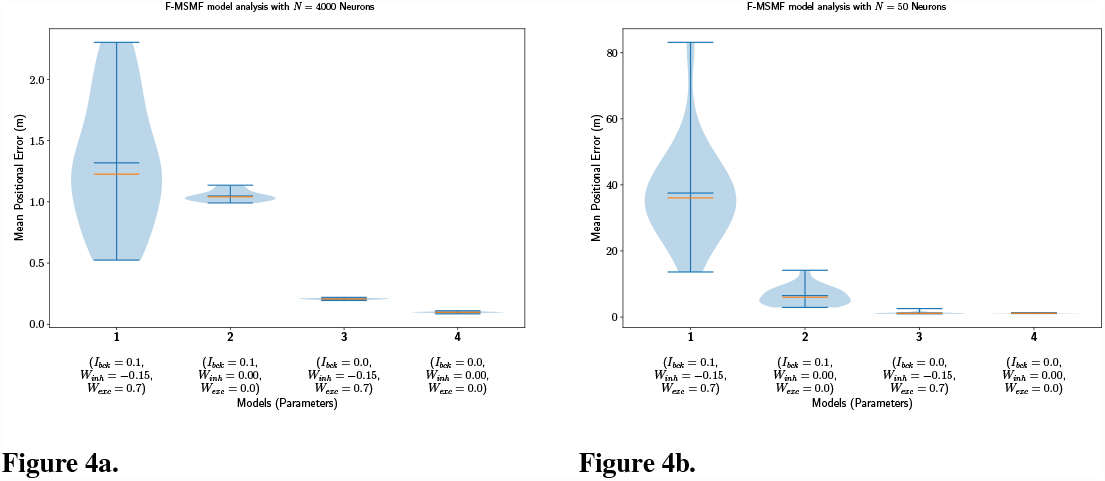
The distribution of the mean positional error of 20 individual runs of the original F-MSMF model with *N* = 4000 neurons (a) as in the results from Eliav et al. (2021a) and *N* = 50 neurons (b) as in the theoretical analysis. The blue lines represent the minimum, maximum, and mean of the evaluation results, the orange line represents the median of it.

In a second experiment, we evaluated the D-MSMF model, introduced in Section 2.1.4. The purpose of this experiment is to create a baseline comparison to the theoretical results by Eliav et al. and also evaluate the network in order to define further experiments for analyzing its properties and performance. We chose the connection parameter *TH*_*fr*_ to be equal to 90%. In the next subsection we will perform a more thorough analysis of this parameter in order to identify more optimal values. The remaining parameters, such as for the gamma distribution of the field sizes, were chosen to be the same as for the theoretical analysis by Eliav et al. The results for *N*_*neu*_ = 50 neurons are visualized in Fig. 5. Interestingly, the median of the average decoded positional error 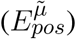 in this case is higher when the lateral connections are removed while the background input persists (model 1 vs. 2). This stands in contrast to the results obtained with the F-MSMF model and might be an indication, that these connections stabilize and denoise the system. Even when comparing the two last runs with each other, although the median and mean error is lower when all lateral connections are removed, the minimum error is even smaller for the third compared to the fourth model (0.858*m* vs. 0.866*m*). The implications of these insights on the significance of lateral connections in MSMF networks are further analyzed in Section 3.4.3.

**Figure 5.**
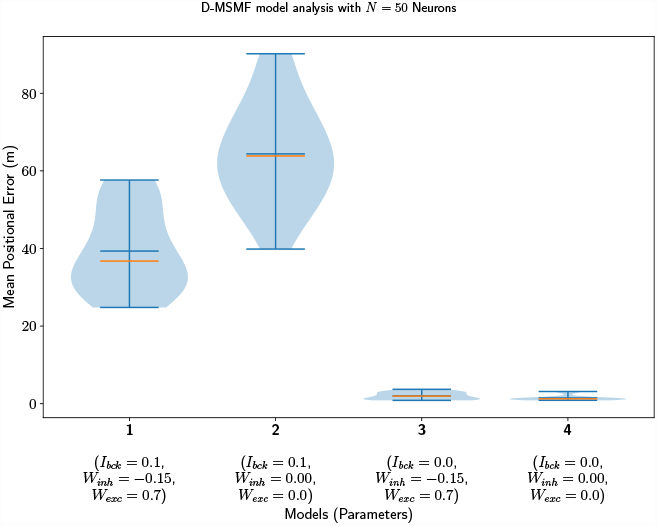
The distribution of the mean positional error of 20 individual runs of the D-MSMF model with *N* = 50 neurons. The blue lines represent the minimum, maximum, and mean of the evaluation results, the orange line represents the median of it.

The results presented in this section show, that, theoretically, the introduced MSMF networks are capable of reproducing the results from the theoretical analysis of Eliav et al. - but only under certain circumstances. The crucial factors that influence the positional encoding performance of these networks are the lateral connections and the noise (background input). In the remainder of this evaluation we will therefore focus not only on the potential theoretical performance of MSMF networks but also on the (dis-)advantages of the lateral connections in such a multi-line attractor as well as the influence of different kinds of noise on the system in order to answer the question, whether a system with such a code could be modeled by an attractor network or not.

### 3.3 MSMF Code

Within this part of the evaluation we focus on the MSMF code itself and therefore only evaluate networks without any lateral connections or background noise, if not stated explicitly:

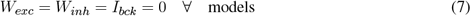

#### 3.3.1 Optimal Parameterization of MSMF Models

The first deeper analysis we perform with the MSMF models is done in order to retrieve the most optimal models with respect to the mean positional error of the network (accuracy). For this evaluation we only optimized the networks for accuracy. We will, however, also compare their expected energy consumption as defined in Section 3.1. The configuration for the evolutionary optimization runs for the F-MSMF as well as the D-MSMF models is shown in Table S1. A visualization of the optimization results can be found in the supplementary materials in Fig. S1a/b and Fig. S2a/b, respectively. For the D-MSMF model, we ran multiple optimizations, continuously shifting the range for *α*, since the results kept improving. We included one row representing all runs - including the average number of generations of all runs.

The sampled parameter combinations for the F-MSMF model, shown in Fig. S1a, illustrate, that in general a higher number of fields, i.e. more neurons per attractor (high *P*_*att*_), is preferable over lower numbers for achieving a low positional decoding error. This completely aligns with the results from the D-MSMF model, visualized in Fig. S2a and S2b. The networks achieving the highest decoding accuracy are all located in the range of *θ <* 0.04. With *θ* this small, the average sampled field size also becomes very small and the number of fields therefore very large, as demonstrated in Fig. S2b, where only networks with a large number of fields are shown. The networks visualized in this figure, correspond to the ones from Fig. S2a with a small positional error.

When further filtering the values of the F-MSMF results (Fig. S1b), one can however see, that at least for this model, a variety of parameter combinations can lead to optimal networks with no positional decoding error 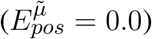. We therefore included three different networks from the optimization results in Table 1. The first two networks achieve an optimal decoding error although the number of fields per neuron differs significantly for each of them. We picked *F-Opt-1* because it maintains the largest number of fields of all optimal network configurations 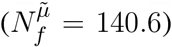 and *F-Opt-2* because it maintains the lowest number of fields 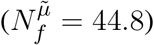 while still having somewhat different scales, i.e. differences between the number of attractors in each level. The third network, *F-Opt-3*, was picked for further analysis in the next sections, as it maintains a low positional error 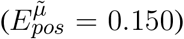 with only 12 fields per neuron 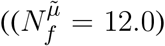. The energy consumption of *F-Opt-2/3* is significantly better than that of *F-Opt-1*, since fields of the neurons cover less space. While this energy consumption is slightly higher, it is significantly smaller than that of *F-Org-1*, showing that a better positional accuracy can be achieved with many small fields while reducing the energy consumption.

**Table 1.** Optimized F-MSMF models without lateral connections.

| Model ID | $N_{AL_0}$ | $N_{AL_1}$ | $N_{AL_2}$ | $P_{att}$ | $N_f^{\tilde{\mu}}$ | $C_{eng}^{\tilde{\mu}}$ | $E_{pos}^{\tilde{\mu}}$ | $E_{pos}^{min}$ | $E_{pos}^{max}$ |
| --- | --- | --- | --- | --- | --- | --- | --- | --- | --- |
| F-Opt-1 | 50 | 48 | 50 | 0.95 | 140.6 | 140.8 | 0.000 | 0.000 | 0.003 |
| F-Opt-2 | 50 | 22 | 40 | 0.40 | 44.8 | 60.0 | 0.000 | 0.000 | 0.004 |
| F-Opt-3 | 11 | 10 | 9 | 0.40 | 12.0 | 59.6 | 0.150 | 0.133 | 0.279 |
| F-Org-1 | 5 | 2 | 1 | 0.30 | 2.4 | 3460.6 | 0.098 | 0.085 | 0.110 |
| F-Org-2 | 5 | 2 | 1 | 0.30 | 2.4 | 43.8 | 1.148 | 1.056 | 1.293 |

This hypothesis is confirmed by the results of the D-MSMF model. As stated before, the optimal values here are all in a range favouring a field size distribution with a small mean and especially a very small variance. For all evaluated networks with a median error 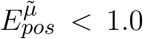, the median of the distribution of all field size means is 0.41, the median of the variance is 0.01. This shows, that for this model the optimal distribution of field sizes results in a large number of very small fields with very little variance in the field size. Due to these properties, the resulting networks can not be defined as a multi-scale model anymore. They further do not achieve an accuracy close the F-MSMF models. Table 2 shows, that the energy consumption is much higher for both, the optimal as well as the original model, than the energy consumption of the F-Opt models (89.5 *>>* 44.8), while the median of the positional decoding error is much higher than that of the most optimal F-MSMF model (*F-Opt-1/2*). This shows that the arrangement of the neurons into multiple, differently sized layers and by that creating fields of very different sizes does seem to have a positive influence on the accuracy of the decoded position.

**Table 2.**
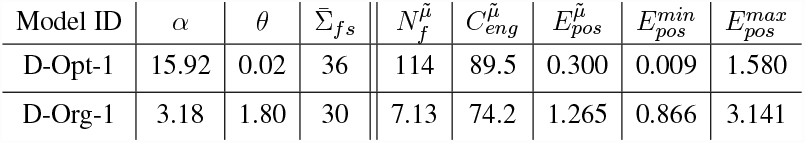
Optimized D-MSMF models without lateral connections.

| Model ID | $\alpha$ | $\theta$ | $\bar{\Sigma}_{fs}$ | $N_f^{\tilde{\mu}}$ | $C_{eng}^{\tilde{\mu}}$ | $E_{pos}^{\tilde{\mu}}$ | $E_{pos}^{min}$ | $E_{pos}^{max}$ |
| --- | --- | --- | --- | --- | --- | --- | --- | --- |
| D-Opt-1 | 15.92 | 0.02 | 36 | 114 | 89.5 | 0.300 | 0.009 | 1.580 |
| D-Org-1 | 3.18 | 1.80 | 30 | 7.13 | 74.2 | 1.265 | 0.866 | 3.141 |

In addition to these findings, especially the D-MSMF models showed a large variance of the decoding error between different runs with the same parameterization but varying initialization of field locations and sizes. For both models, D-Org and D-Opt-1, the discrepancy between the minimum of all mean decoding errors of 20 runs and the maximum is significant 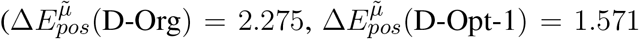. Since it occurs for both models almost at an equal level, the shape of the gamma distribution as well as the number of fields do not seem to have an impact on it. This instability of the networks will be further investigated in the next part of this evaluation.

In order to further investigate the optimal parameterization of the network, we analyzed the influence of the maximal field coverage of a neuron 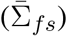. For this experiment, we ran the original D-MSMF model (D-Org-1) and varied the value for 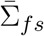 between each run. We chose a range from 1 to 100 here. The median of the resulting positional error is visualized in Fig. 6. This result demonstrates two things. First, the mean/median measured experimentally (30*m*) by Eliav et al. (2021a) lies within the minimum of this plot, which further confirms the parameter and model choice. In the experiments, however, many cells experienced a field coverage much larger than that. In fact, a significant portion of the cells had a field coverage 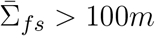. According to our investigation of this model with the given parameters, this would lead to a significant drop of the positional decoding accuracy (*>* 10*m*) compared to the optimal values at the minimum around 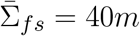. This, on the other hand, is an indicator that either this model, or at least its parameters, are not suited for representing the given MSMF code, or the given MSMF code is not just a “simple” place code as thought for decades.

**Figure 6.**
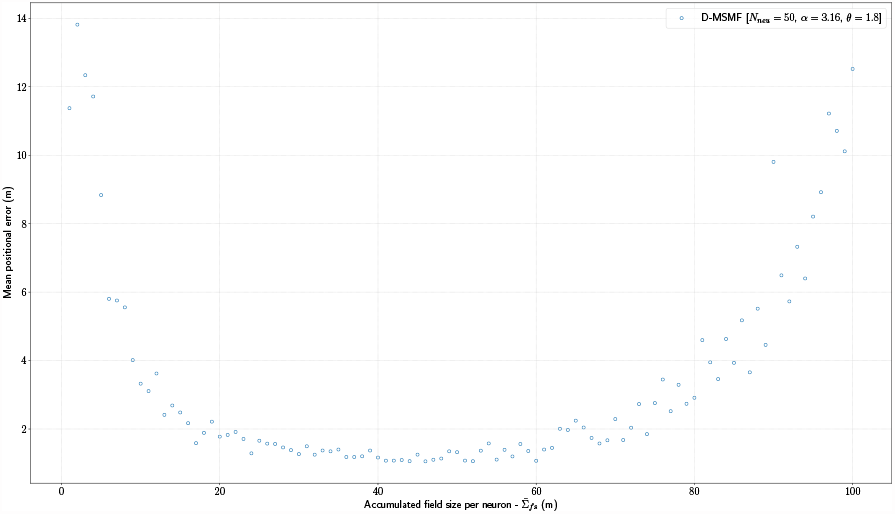
The median positional error for a range of experiments performed with the *D-Org-1* model, varying the maximal field coverage 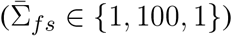.

#### 3.3.2 D-MSMF Variation Analysis

In the previous section, we optimized the MSMF models in order to identify optimal parameter combinations. This experiment also demonstrated, that these networks are highly unstable, i.e. the same parameterization does not necessarily lead to the same or even a similar accuracy. Within this part of our evaluation we therefore evaluated these extreme scenarios in which a network with the same parameters produces a large and a small error when initialized differently. The goal of this evaluation was to identify possible factors of place field distribution which have a (non-)benefitial impact on the decoding accuracy, such as a uniform distribution similar to the grid code, or find some properties which lead to errors in the decoding, such as a high number of falsely active cells.

In order to evaluate this, we compared the results of the *D-Opt-1* and the *D-Org-1* model (see Table 2). Both networks have a high variation between the minimum and maximum mean positional decoding error, which can be reached with the same parameters and different field initializations. They do, however, differ significantly in their field size distribution, hence model *D-Opt-1* has a large number of fields 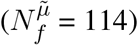 while model *D-Org-1* has a low number of fields 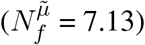.

The analysis we conducted in order to identify possible problems with these networks include

- the percentage of unique field combinations,
- the average number of false positive/negative bins,
- the average distance between all field locations and the nearest bin location (centers),
- the divergence of field size/location distribution from their respective actual distribution.

The results of these analysis are visualized in the supplementary materials. They do not indicate that there is any kind of pattern, convergence or correlation between the decoded positional error 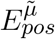 of a network and any of these properties. Our best guess in this case is, that these high variances occur because of the natural drift caused by the dynamical system. This can result in a situation, where a cell is more active in a bin which is not counted as part of its field while another cell which is expected to be active in that bin, since its field is overlapping with it, has a lower activity there. This is a corner case of false positives that we did not analyze as it is non-trivial. In order to investigate this further, one has to analyze false positives not only qualitatively but also quantitatively. We elaborate more on this topic further in the future work.

#### 3.3.3 Benchmark with Grid Code

In order to put the results from the original and optimized MSMF models into context, we compare them in this section to the results from multiple optimized one-dimensional grid codes. Each code is build by a network with multiple modules (*N*_*mod*_), each of which contains a certain number of neurons 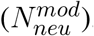. The modules have different scales, with a minimum scale 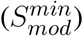 and a multiplier from one scale to the next (*S*_*mod*_). All these parameters were optimized over 3000 epochs without any lateral connections. The results of a few exemplary networks with optimal positional decoding error but different properties are visualized in Table 3.

**Table 3.** Optimized grid models without lateral connections.

| Model ID | $N_{mod}$ | $N_{neu}^{mod}$ | $S_{mod}$ | $S_{min}^{mod}$ | $N_f^{\tilde{\mu}}$ | $C_{eng}^{\tilde{\mu}}$ | $E_{pos}^{\tilde{\mu}}$ | $E_{pos}^{min}$ | $E_{pos}^{max}$ |
| --- | --- | --- | --- | --- | --- | --- | --- | --- | --- |
| G-Opt-1 | 3 | 9 | 1.6 | 0.5 | 29.852 | 30.75 | 0.0 | 0.0 | 0.0 |
| G-Opt-2 | 9 | 19 | 3.0 | 0.5 | 3.509 | 64.57 | 0.0 | 0.0 | 0.0 |
| G-Opt-3 | 9 | 7 | 3.0 | 0.5 | 9.524 | 64.64 | 0.0 | 0.0 | 0.0 |
| G-Opt-4 | 3 | 19 | 1.2 | 0.5 | 17.737 | 30.76 | 0.0 | 0.0 | 0.0 |
| G-Opt-5 | 3 | 19 | 1.8 | 0.5 | 13.07 | 30.73 | 0.0 | 0.0 | 0.0 |

The optimization of the grid code shows that with at least 3 modules and 4 neurons per module, almost all combinations of the grid model achieve the same or even better positional decoding accuracy as the best optimized MSMF models introduced so far. We picked five samples from the optimized models with different number of modules, neurons per module and module scale, all of them achieving a median decoding error of 0*m*. The networks can be categorized as follows:

**G-Opt-1** Lowest number of neurons overall (27).

**G-Opt-2** Largest number of neurons overall (171).

**G-Opt-3** Large number of modules and a small number of neurons. **G-Opt-4** Large number of neurons and a small number of modules. **G-Opt-5** Same as *G-Opt-4* but with a much larger module scale.

The reason why we picked these models is to evaluate the performance of different combinations of module size, number of neurons and module scale. In the evaluation results focussing on the positional decoding error and energy consumption, shown in Table 3, there are no differences in the accuracy between the networks. The energy consumption, on the other hand, increases significantly when the number of modules rises. This can be expected, since each new module adds another layer of 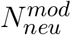 neurons, resulting in additional activity and hence an increased energy consumption.

In order to analyze the robustness of all models described so far in this evaluation, we conducted further experiments with a certain percentage of drop-out neurons. Fig. 7 visualizes the results for this experiment. By far the best performing model is, as expected, the *F-Org-1* with 4000 neurons overall. Even in the worst case, with 95 % of the neurons being dead, it still performs better than most networks with just 5 % lesions. All of the optimized F-MSMF models (*F-Opt-1/2/3*) are capable of maintaining a median positional decoding error 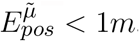, even with a drop-out rate of *P*_*dro*_ = 0.25, i.e. 25 % randomly removed neurons. This demonstrates the effect of the redundancy in these models, caused by the large number of fields per neuron. This redundancy and robustness seems to be more efficient for *F-Opt-2/3* than the mechanism that is needed in the grid code. Almost all of the models here perform significantly worse than the rest, even with only 5 % of the neurons being disabled. Only the *G-Opt-2* model performs comparably well to the optimized F-MSMF models. It does, on the other hand, require a significantly larger amount of neurons for that 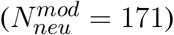. This shows, that in order to gain robustness in grid models, one needs a large number of modules and neurons to achieve redundancy.

**Figure 7.**
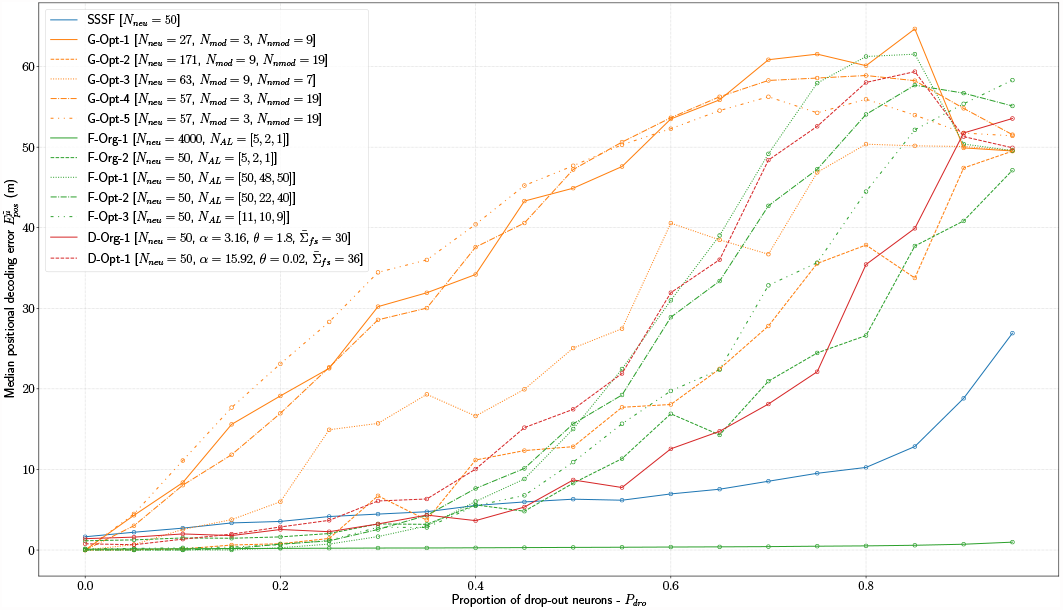
Evaluation of all introduced models (F-MSMF, D-MSMF, Grid, SSSF) with an increasing percentage of drop-out neurons (*P*_*dro*_ ∈ {0.0, 0.95, 0.05}).

### 3.4 Lateral Connections in MSMF Models

The last part of our evaluation focuses on the lateral connections in the MSMF models, i.e. the connections which are essential for making it a CAN.

#### 3.4.1 Optimized MSMF Models with Lateral Connections

For the proper evaluation of the purpose or benefits of the lateral connections in the MSMF models we performed multiple optimizations of the models with different parameterizations. For each model (D-MSMF, F-MSMF) we performed three optimizations: the first one optimizes for all parameters of the network (lateral connections and architecture/field distribution), the other two only for the lateral connections while the architecture and field distribution remain constant (original and optimal parameters). The parameters for training the networks are listed in Table S4, the trained parameters of the networks are listed in Tables S5 and S6. The optimization results are visualized in Fig. 8.

**Figure 8.**
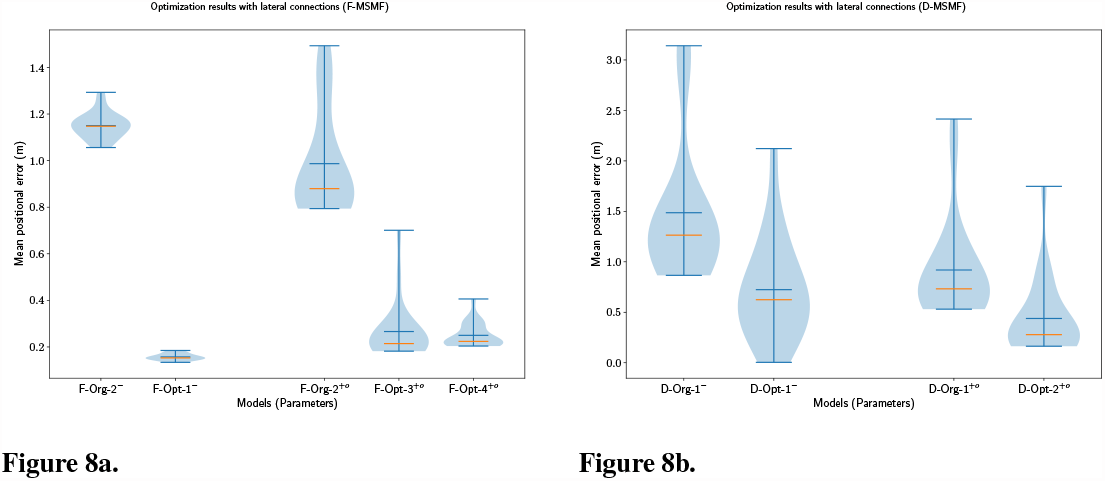
The distribution of the mean positional error of 20 individual runs for optimized F-MSMF (a) and D-MSMF (b) models with lateral connections. On the left of each figure the results from the previous experiments without lateral connections are shown. On the right, the results from the optimization are visualized. The blue lines represent the minimum, maximum, and mean of the evaluation results, the orange line represents the median of it.

For the F-MSMF model, we optimized the lateral connection parameters for the *F-Org-1* and *F-Opt-3* models, resulting in the *F-Org-1*^+*o*^ and *F-Opt-3*^+*o*^ models, respectively. An initial optimization of the *F-Opt-1/2* models with lateral connections resulted in positional decoding errors far to high for further experiments, even after several hundred epochs of training. We therefore picked the *F-Opt-3* model, as it led to a reasonably low positional decoding error with optimized lateral connections. In addition to that we optimized all parameters, including the architectural parameters, resulting in the new model *F-Opt-4*^+*o*^. We kept the maximum number of attractors per level quite low in this case, due to the aforementioned issue with training lateral connection weights for models with a large number of attractors (*N*_*AL*_ *>>* 30).

The evaluations of these networks (Fig. 8a) show, that the lateral connections reduce the median decoding error for the original network architecture (*F-Org-2*^*−*^ vs. *F-Org-2*^+^) and increase it for the optimized architecture (*F-Opt-3*^*−*^ vs. *F-Opt-3*^+*o*^/*F-Opt-4*^+*o*^). This indicates that lateral connections are more beneficial in a spatial code with fewer but larger fields per neuron, since the *F-Opt-3* model has significantly more fields per neuron than the *F-Org-2* model 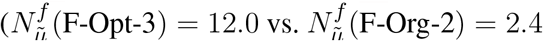, cmp. Table 1).

We performed the same three optimizations for the D-MSMF model. The results shown in Fig. 8b do not depict the results for the optimization of the *D-Opt-1* model, however. The reason for that is that this optimization did not lead to any results. After running it for 200 generations, the median decoding error was still around 50 m. We therefore omitted this result. This result together with those of the remaining two optimizations confirm the indications that the analysis of the F-MSMF optimizations already revealed - especially networks with fewer and larger fields benefit from lateral connections. This seems intuitive, since more fields also lead to more connections and with that to more noise. Creating only a few connections with small weights, however, seems to stabilize the system and reduce noise.

In addition to that, we observed that most of the weights of the optimized models were in fact negative, for some of them even all weights. This applied especially to the cases where the decoding error dropped by introducing the optimized weights. We will analyze the influence of the weights on the firing fields of individual neurons further in Section 3.4.3.

In order to analyze the general benefit of connecting two neurons based on the individual field sizes of these neurons, we performed an experiment with D-MSMF models. In this experiment two different models were chosen (*D-Org-1*^+^ and *D-Opt-3*^+^) with lateral connections according to the previous optimizations results. Both models were evaluated 100 times, one time with a field ratio threshold (*TH*_*fs*_(*D-Org-1*^+^) = 0.83 and *TH*_*fs*_(*D-Opt-3*^+^) = 0.79) and a field connection probability (*P*_*fc*_(*D-Org-1*^+^) = 0.87 and *P*_*fc*_(*D-Opt-3*^+^) = 0.76). The results of these experiments are visualized in Fig. 9. These results demonstrate that there is no benefit in creating connections between neurons based on their respective field sizes. Creating random connections leads to very similar but in both cases here even smaller decoding errors. While we do not have an explanation for the decrease of the decoding error, we observed, that the fields of the networks with a field connection probability were sharpened equivalently to the sharpening which occurs when using a field ratio threshold.

**Figure 9.**
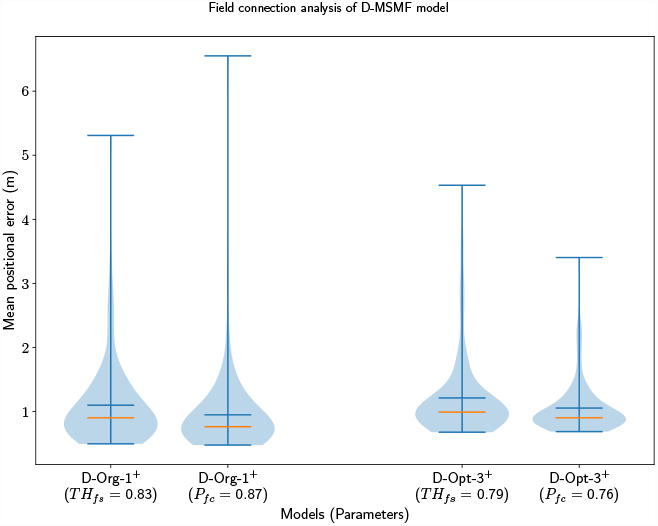
The distribution of the mean positional error of 100 individual runs of pairs of D-MSMF models, with either the field ratio threshold or an equivalent field connection probability set. The blue lines represent the minimum, maximum, and mean of the evaluation results, the orange line represents the median of it.

#### 3.4.2 CAN Features in MSMF Models

One of the key features of CANs is the maintenance of a bump of activity in the absence of a specific input. Some networks are capable of maintaining a bump of activity after the specific input is removed without receiving any kind of input at all, while others need a certain amount of unified background input, all depending on the setup of the connections between neurons. In this part of the evaluation we have locked at both of these cases for evaluating whether the MSMF models can achieve this and are indeed Continuous Attractor Networks or not. For this purpose we create a baseline with an SSSF model with *N*_*neu*_ = 50 neurons spanning uniformly over the whole environment. We then remove the input for a length of *L*_*rem*_ = 20*m* and evaluate the network with and without lateral connections. If the lateral connections do create a CAN, then the decoding error is expected to be smaller with lateral connections present. During the time, where the positional input is removed, the optimal decoded position is standing still, i.e. it is equal to the last position where the positional input was active. This leads to a scenario, where the maintenance of a bump at the last known location after the positional input is removed leads to an optimal decoded position.

We picked multiple different models from the previous experiments and optimizations in order to verify, that the results are not based on a certain parameterization of the network. For the F-MSMF model, we chose the *F-Opt-3* as a reference, since we could not get any of the other networks optimized with lateral connections (see Section 3.4.1). The decoded error for all experiments is shown in Fig. 10. The models are visualized pairwise, without and then with lateral connections. If the respective model is a CAN, then the error should decrease from the first to the second run, as it is the case for the SSSF model (*S-Std-1*). This does, however, not apply for any of the MSMF models. On the contrary, the error increases significantly for all of the MSMF models. These results show, that there is no evidence that these MSMF models are indeed CANs.

**Figure 10.**
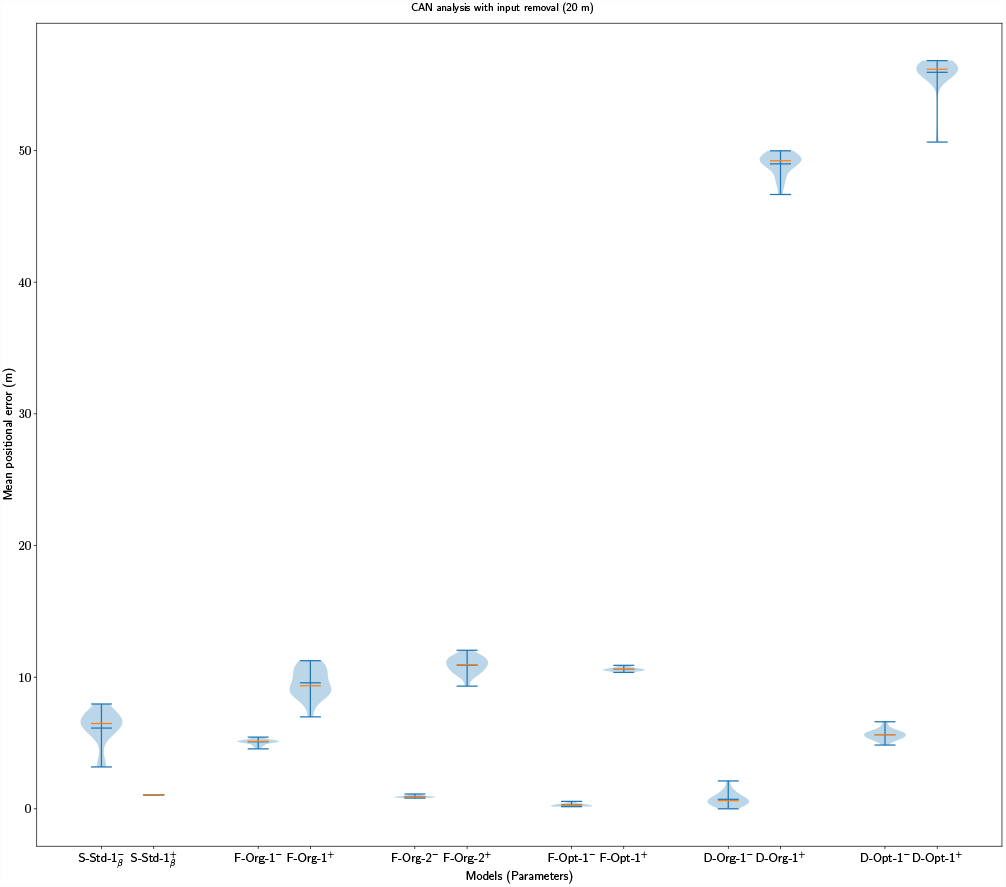
The distribution of the mean positional error of 20 individual runs of pairs of SSSF, F-MSMF and D-MSMF models (without/with lateral connections). The blue lines represent the minimum, maximum, and median of the evaluation results, the orange line represents the mean of it.

#### 3.4.3 Benefits of Lateral Connections in MSMF Models

In the previous part of the evaluation, we have shown that the MSMF models do not seem to fulfill some typical properties of a CAN. In this final part of the evaluation we now investigate what other benefits or purpose the lateral connections could have in such a model. We therefore analyze the influence of the lateral connections on the field shape of the individual neurons in both the F- and D-MSMF models.

The results of this analysis are visualized in Fig. 11 and Fig. 12 for the F-Org-2^*−/*+^ and D-Org-1^*−/*+^ models, respectively. In both cases, the activation of the lateral connections leads to a sharpening of almost all firing fields. Due to this sharpening the fields have less activity outside of their actual field and hence lead to less noise in the decoding (false positives). At least for the D-Org-1^*−/*+^ model we already showed in 3.4.1, that enabling the lateral connections leads to a decrease of the median positional decoding error.

**Figure 11.**
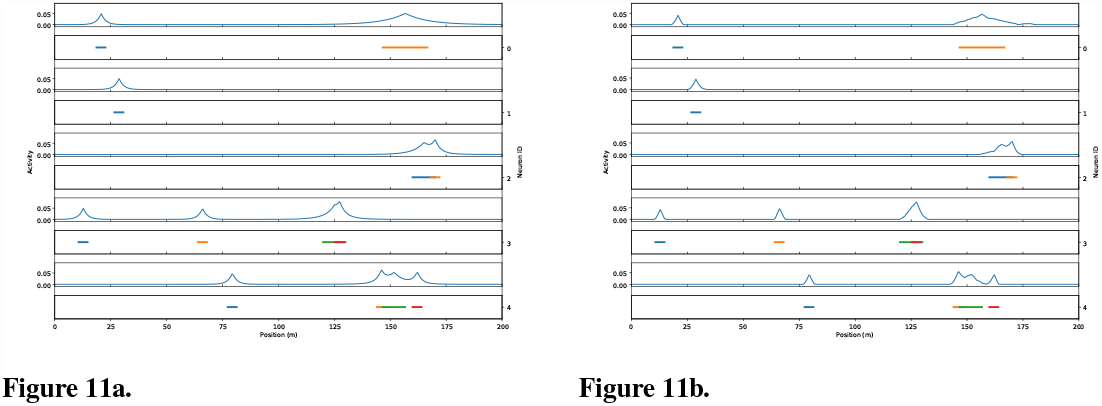
The field activity for the first five neurons of the first network of the experiment performed with 20 instances of F-Org-2^*−*^ (a) and F-Org-2^+^ (b).

**Figure 12.**
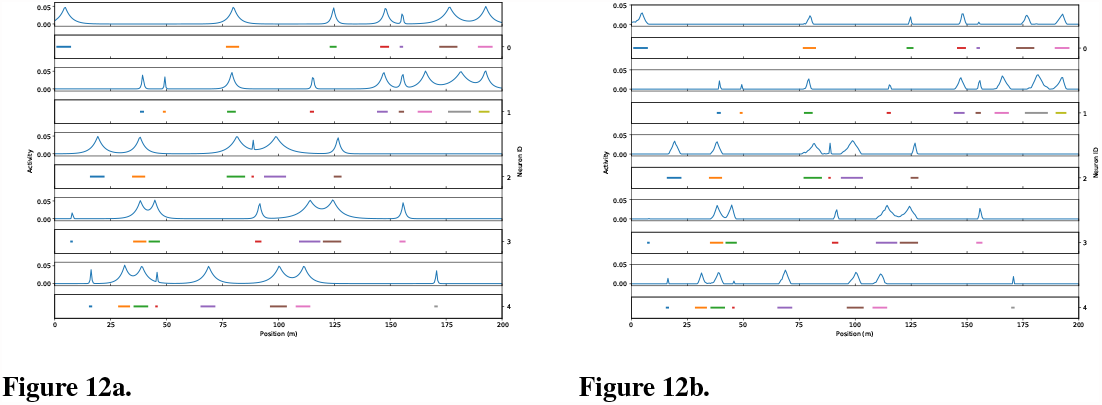
The field activity for the first five neurons of the first network of the experiment performed with 20 instances of D-Org-1^*−*^ (a) and D-Org-1^+^ (b).

Although this does not apply to all of the models, we do think that lateral connections in such a model could be used for de-noising the input data. This, however, seems to require few connections with small negative weights. We modified the field size threshold *TH*_*fs*_ for creating connections between two neurons and set it to *TH*_*fs*_ = 0.8 for the D-Org-1^+^ model. This lead to a shift of the weight distribution to much smaller values. While the same number of connections was established, the weight of each connection was significantly smaller than before. The median decoding error dropped as well to 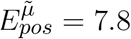. When analyzing the field distributions of the individual neurons in that model, we realized that the fields of each neuron have shrunken even more, hence leading to many false-negatives during the decoding (1881 vs. 970).

## 4 CONCLUSION

This paper aimed to evaluate the accuracy and robustness of a recently found multi-scale multi-field place code in the hippocampus of bats, together with possible CAN-based network topologies producing such a code. In order to achieve that, we trained several networks using evolutionary optimization and compared the MSMF networks to an SSSF network (line attractor) as well as a grid code.

Based on this analysis, two main contributions were presented in this paper. First, we identified that, although the MSMF code does outperform an SSSF code with respect to the decoding accuracy, it is outperformed by most grid codes we explored. Like the MSMF codes, grid codes also contain fields of different sizes, but distribute these fields optimally in environments of any dimension Mathis et al. (2015). Our experiments on MSMF codes have shown that even with the same parameters for field generation, the decoding accuracy can change significantly between multiple instances of the same network with different field locations and combinations. Due to the much larger number of fields in some of the MSMF models, they are, however, much more robust to noise induced by drop-out or lesions. This can be explained by the redundancy introduced by the large number of fields per neuron compared to the grid code.

The second contribution in this paper is our analysis of the multi line attractor networks that were first proposed by Eliav et al. (2021a). We showed that these networks do not exhibit common properties of continuous attractors. When removing the position-dependent input for a short period of time, the network would always converge to a single baseline attractor state, independently of where the animal currently is. During the movement of the agent/animal, this discrete attractor is always active in the background. It is, however, overpowered by the location specific input to the network, as long as this is provided. The only benefit of the lateral connections we could identify in these networks was their ability to create more precise firing fields by introducing inhibition, which trims the “foothills” of the activity bumps.

Based on these results, we conclude that the MSMF place code found in the hippocampus of bats is not the most suitable representation with respect to accuracy and energy efficiency, unless robustness to noise is also taken into account. Surprisingly, we found that the MSMF codes we investigated did not have continuous attractors. It is, therefore, possible that the bats’ MSMF code does not originate from a continuous attractor network topology.

## ACKNOWLEDGMENTS

We thank Misha Tsodyks for providing us with the original code of the multi-scale, multi-field network, Andreas Herz for his support and Benjamin Dunn for his feedback and discussions.

## DATA AVAILABILITY STATEMENT

The framework we developed within this study and used for all the simulations, optimizations and experiments can be found at https://github.com/dietriro/msmf-code. It contains all models as well as evaluations and is partially based on the code by Eliav et al. (2021a).

