## Supplementary Materials for "Grid Codes versus Multi-Scale, Multi-Field Place Codes for Space"

### 1 SUPPLEMENTARY MATERIALS

#### 1.1 Evaluation

##### 1.1.1 Optimal Parameterization of MSMF Models

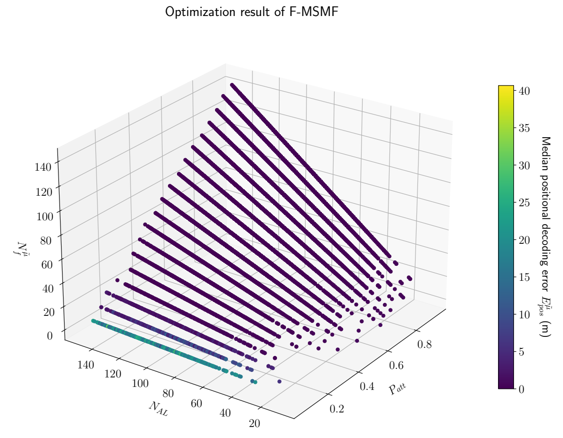


**Figure S1a**

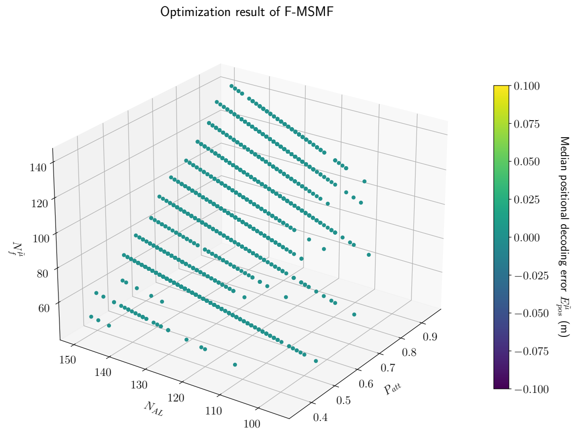


**Figure S1b**

**Figure S1.** Results of F-MSMF optimization after 3000 generations. Visualized on the axis are the overall number of attractors at all levels ($N_{AL}$), the probability of participation in an attractor ($P_{att}$) and the median number of fields ($N_f^{\tilde{\mu}}$). The colorbar shows the median positional decoding error ($E_{pos}^{\tilde{\mu}}$). The results are filtered by the median positional error: **a)** ($E_{pos}^{\tilde{\mu}} < \infty$), **b)** ($E_{pos}^{\tilde{\mu}} == 0.0$)

##### 1.1.2 Lateral Connections in MSMF Models

##### 1.1.3 D-MSMF Field Size/Location Distribution

In the previous section, we optimized the MSMF models in order to identify optimal parameter combinations. This experiment also demonstrated, that these networks are highly unstable, i.e. the same parameterization does not necessarily lead to the same or even a similar decoded positional error. Within this part of our evaluation we therefore evaluate specifically these extreme scenarios, in which a network with the same parameters produces a large and a small error when initialized differently. The goal of this evaluation is to identify possible factors of place field distribution which have a (non-)benefitial impact on the decoding accuracy, such as a uniform distribution similar to the grid code.

In order to evaluate this, we picked two models from the optimization results in the previous paragraph. Both networks have a high variation between the minimum and maximum mean positional decoding error which can be reached with the same parameters and different field initialization. We chose one model (A) with a large number of fields $N_f^{\mu}$ (high MAFS) and another model (B) with a low number of fields $N_f^{\mu}$ (low MAFS). The parameters for both models are shown in Table S5 together with some evaluation results on population level.

**Table S1.** Parameters for optimization of F-/D-MSMF models without lateral connections.

| Netw. Type | #Gen. | Param 1 | Param 2 | Param 3 | Param 4 |
|---|---|---|---|---|---|
| F-MSMF | 3000 | $N_{AL_0} \in \{1, 50, 1\}$ | $N_{AL_1} \in \{1, 50, 1\}$ | $N_{AL_2} \in \{1, 50, 1\}$ | $P_{att} \in \{0.05, 1.0, 0.05\}$ |
| D-MSMF | 1610 | $\alpha \in \{0.02, 30.00, 0.02\}$ | $\theta \in \{0.02, 6.00, 0.02\}$ | $\bar{\Sigma}_{fs} \in \{2, 100, 1\}$ | |

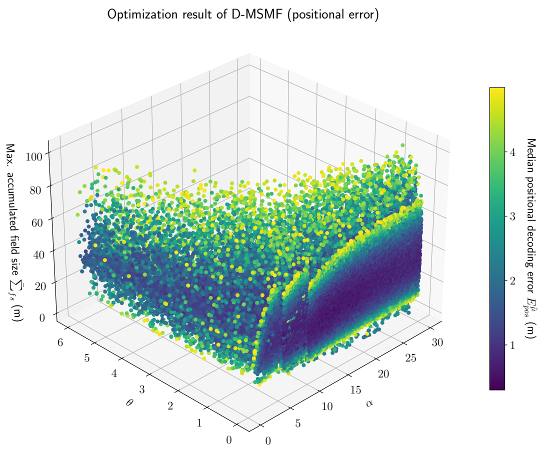


**Figure S2a**

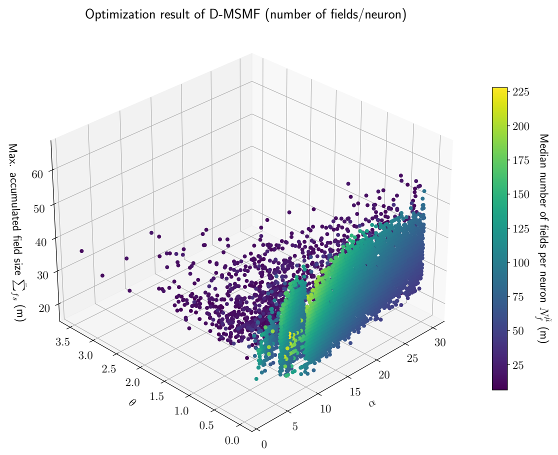


**Figure S2b**

**Figure S2.** Results of D-MSMF optimizations after 1600 generations. Visualized are the parameters of the network on the axis ($\alpha$, $\theta$ and $\bar{\Sigma}_{fs}$) and on the colorbar the median positional error (**a**)) as well as the median number of fields per neuron (**b**)). The results are filtered by the median positional error: **a)** ($E^{\tilde{\mu}}_{pos} < 5.0$), **b)** ($E^{\tilde{\mu}}_{pos} < 1.0$)

We then calculated the Kullback-Leibler divergence (D) for the field size distribution (with a gamma distribution) and the field location distribution of each neuron per network. The mean ($D^{\mu}$) and standard deviation ($D^{\sigma}$) of all networks is visualized in Figures S3a and S3b for the field sizes and locations, respectively.

The results shown in these figures do not indicate that there is any kind of pattern, convergence or correlation between the decoded positional error $E^{\tilde{\mu}}_{pos}$ of a network and its field location/size initialization. On the contrary, it seems like all networks, independently from their decoding accuracy, are similarly close to their respective true distribution. The small, overall divergence of the networks can be explained by the way the fields sizes and locations are initialized. The maximal accumulative field size (MAFS) restricts the number and sizes of the fields, especially when it is almost reached. We then try to find a size, based on the gamma distribution, which would still fit without exceeding the MAFS. We repeat this process for maximally $N_{try}$ times, hence the actual field size distribution can diverge slightly from the original one. Since fields are not allowed to overlap, it can also happen, that the location has to be sampled many times before one without overlap is found. This can negatively impact the KL divergence from the uniform distribution.

**Table S2.** Parameters for optimization of F-/D-MSMF models with lateral connections.

| Netw. Type | Param 1 | Param 2 | Param 3 | Param 4 | Param 5 | Param 6 |
|---|---|---|---|---|---|---|
| F-MSMF | $N_{AL_0} \in \{1, 11, 1\}$ | $N_{AL_1} \in \{1, 11, 1\}$ | $N_{AL_2} \in \{1, 11, 1\}$ | $P_{att} \in \{0.1, 0.5, 0.05\}$ | $W_{exc} \in \{0.02, 1.6, 0.02\}$ | $W_{inh} \in \{-0.08, -0.02, 0.02\}$ |
| D-MSMF | $\alpha \in \{0.02, 30.00, 0.02\}$ | $\theta \in \{0.02, 6.00, 0.02\}$ | $\bar{\Sigma}_{fs} \in \{2, 100, 1\}$ | | $W_{exc} \in \{0.02, 1.6, 0.02\}$ | $W_{inh} \in \{-0.08, -0.02, 0.02\}$ |

**Table S3.** Optimized F-MSMF models with lateral connections. Only bold values were trained.

| Model ID | $W_{inh}$ | $W_{exc}$ | $I_{bck}$ | $N_{AL_0}$ | $N_{AL_1}$ | $N_{AL_2}$ | $P_{att}$ | $N_f^{\tilde{\mu}}$ | $C_{eng}^{\tilde{\mu}}$ | $E_{pos}^{\tilde{\mu}}$ | $E_{pos}^{min}$ | $E_{pos}^{max}$ |
|---|---|---|---|---|---|---|---|---|---|---|---|---|
| F-Org-2$^{+o}$ | **-0.1** | **0.04** | 0.0 | 5 | 2 | 1 | 0.3 | 2.4 | 27.08 | 0.880 | 0.794 | 1.493 |
| F-Opt-1$^{+o}$ | **-0.04** | **0.2** | 0.0 | 11 | 10 | 9 | 0.4 | 12.0 | 25.46 | 0.215 | 0.182 | 0.701 |
| F-Opt-4$^{+o}$ | **-0.04** | **0.22** | 0.0 | **11** | **10** | **9** | **0.45** | 13.5 | 27.079 | 0.224 | 0.204 | 0.406 |

**Table S4.** Optimized D-MSMF models with lateral connections. Only bold values were trained.

| Model ID | $W_{inh}$ | $W_{exc}$ | $I_{bck}$ | $TH_{fs}$ | $\alpha$ | $\theta$ | $\bar{\Sigma}_{fs}$ | $N_f^{\tilde{\mu}}$ | $C_{eng}^{\tilde{\mu}}$ | $E_{pos}^{\tilde{\mu}}$ | $E_{pos}^{min}$ | $E_{pos}^{max}$ |
|---|---|---|---|---|---|---|---|---|---|---|---|---|
| D-Org-1$^{+o}$ | **-0.04** | **0.08** | 0.0 | **0.83** | 3.16 | 1.8 | 30 | 7.03 | 27.609 | 0.733 | 0.531 | 2.415 |
| D-Opt-2$^{+o}$ | **-0.04** | **0.58** | 0.0 | **0.99** | **2.7** | **0.72** | **40** | 23.35 | 41.042 | 0.279 | 0.164 | 1.748 |

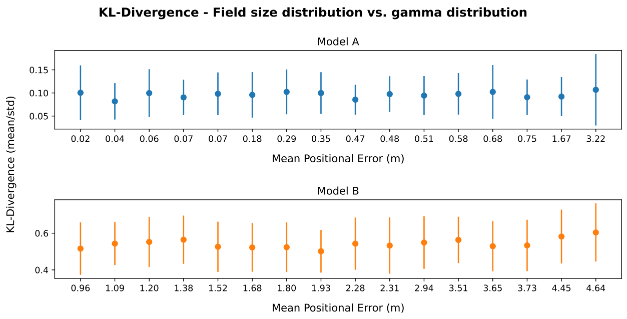


**Figure S3a**

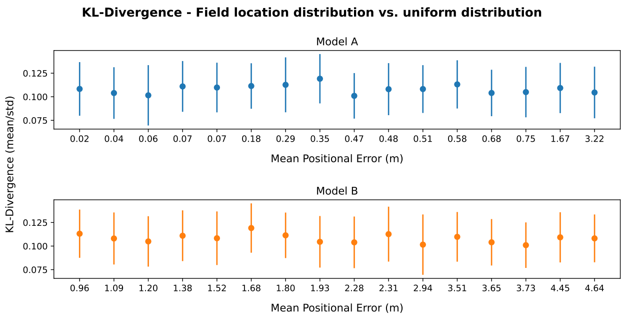


**Figure S3b**

**Figure S3.** The mean positional error of the original F-MSMF model with $N = 4000$ neurons (a) as in the results from Eliav et al. (2021) and $N = 50$ neurons (b) as in the theoretical analysis.

**Table S5.** Evaluated models.

| ID | Network Type | $\alpha$ | $\theta$ | $\bar{\Sigma}_{fs}$ | $N_r$ | $N_f^\mu$ | $\bar{E}_p^\mu$ | $\bar{E}_p^{min}$ | $\bar{E}_p^{max}$ |
|---|---|---|---|---|---|---|---|---|---|
| A | D-MSMF | 3.18 | 1.8 | 30 | 100 | 7.0 | 1.31 | 0.866 | 4.81 |
| B | D-MSMF | 15.92 | 0.02 | 36 | 100 | 114.0 | 0.63 | 0.004 | 1.95 |

The higher divergence of the field sizes of model B from the gamma distribution ($\alpha = 2.02, \theta = 1.8$) can be explained by the combination of a small MAFS with a gamma distribution that returns fields larger than 2.0 m in $70\%$ of the cases. This can result in an overproportional number of small fields, relative to the original (gamma) distribution.

Overall we could not find any hint or reason, why a network with the same parameters can have such a high variance in its decoded positional accuracy. In order to investigate this further, one has to analyze the exact distribution of the field locations and their sizes, i.e. find patterns, groups, or other irregularities. We elaborate more on this topic further in the future work.

**Table S6.** Network and optimization parameters, as well as metrics.

| Parameter | Description | Default value |
|---|---|---|
| **Global parameters** | | |
| Network | | |
| $N_{neu}$ | Number of neurons | $50$ |
| $I_{bck}$ | Uniform background input | $0.1$ |
| $I_{loc}$ | Location specific input | $0.05$ |
| $W_{exc}$ | Maximum excitatory weight | $0.7$ |
| $W_{inh}$ | Minimum inhibitory weight | $-0.15$ |
| $P_{dro}$ | Percentage of drop-out (dead) neurons | – |
| Simulation | | |

| | | |
|---|---|---|
| $T$ | Total simulation time (s) | 20 |
| $\tau$ | Time co-efficient | 0.01 |
| Environment | | |
| $L_{env}$ | Environment length (m) | 200 |
| $\delta_{dsc}$ | Discretization step (m) | 0.5 |
| $N_{bins}$ | Number of bins in the environment | 400 |
| **F-MSMF specific parameters** | | |
| $P_{att}$ | Probability of attractor participation for one neuron | 0.3 |
| $N_{AL_0}$ | Number of attractors in minimum layer | 1 |
| $N_{AL_1}$ | Number of attractors in medium layer | 2 |
| $N_{AL_2}$ | Number of attractors in maximum layer | 5 |
| $L_{int}$ | Interaction length between neurons in one attractor | 5% |
| $sample_{replace}$ | Sample neurons for attractor w/o replacement | $False$ |
| **D-MSMF specific parameters** | | |
| $\alpha$ | Alpha of gamma-distribution | 3.16 |
| $\theta$ | Theta of gamma-distribution | – |
| $TH_{fs}$ | Threshold of the field size ratio | – |
| $P_{fc}$ | Field connection probability | – |
| $\bar{\Sigma}_{fs}$ | Maximum allowed sum of all field sizes of one neuron (m) | 30 |
| **Grid model specific parameters** | | |
| $N_{mod}$ | Number of modules | – |
| $N_{neu}^{mod}$ | Number of neurons per module | – |
| $S_{mod}$ | Scale of the modules | – |
| $S_{mod}^{min}$ | First/minimal scale of all modules | – |
| **Metrics** | | |
| $N_f^{\tilde{\mu}}$ | Median number of fields per neuron | – |
| $E_{pos}^{\tilde{\mu}}$ | Median error of multiple runs | – |
